# Altered Dorsolateral Prefrontal Glutamate Dynamics During Working Memory in Trauma-Exposed Individuals With and Without PTSD: A 7T Functional Magnetic Resonance Spectroscopy Study

**DOI:** 10.64898/2026.08.10.744040

**Authors:** Alex S. Beaver, Sarah E. Whiteman Sitts, Abigail A. Camden, Stephanie M. Jeffirs, Frank W. Weathers, Thomas S. Denney, Meredith A. Reid

## Abstract

Post-traumatic stress disorder (PTSD) has been associated with impairments in cognitive function, including working memory, and may involve altered glutamatergic regulation in the prefrontal cortex. In this study, we used 7T functional magnetic resonance spectroscopy (fMRS) to examine dorsolateral prefrontal cortex (DLPFC) glutamate during working memory in individuals with PTSD, trauma exposure without PTSD (TE), and no trauma exposure (NT). Eighty participants (27 PTSD, 27 TE, 26 NT) underwent baseline MRS followed by fMRS during a letter n-back task. A linear mixed-effects model was used to evaluate glutamate concentrations across baseline, 0-back, 1-back, 2-back, and post-task fixation conditions. Behavioral performance was assessed using repeated-measures ANOVA for percentage correct, reaction time, and the discrimination index (*d’*) across the 0-back, 1-back, and 2-back conditions. Glutamate differed significantly by group, condition, and the group × condition interaction. Individuals with PTSD exhibited lower glutamate than NT at baseline and during the 0-back, 1-back, and 2-back conditions. TE participants also showed lower glutamate than NT during the 1-back and 2-back conditions. Within-group analyses showed higher glutamate during the 0-back, 1-back, and 2-back conditions than at baseline in the NT group, whereas these baseline-to-task differences were limited in the PTSD and TE groups. Accuracy decreased and reaction time increased with increasing working memory load, and discrimination (*d’*) was lower in PTSD than NT. These findings demonstrate altered DLPFC glutamate dynamics during working memory in PTSD and trauma-exposed individuals. Functional MRS provides complementary information beyond resting-state MRS by characterizing glutamatergic responses during cognitive engagement and may improve our understanding of neurochemical alterations associated with trauma and PTSD.

## 1. INTRODUCTION

Exposure to traumatic events can lead to post-traumatic stress disorder (PTSD), a psychiatric disorder with a lifetime prevalence of approximately 8%^1,2^ PTSD is characterized by symptoms of intrusions (e.g., memories, dreams), avoidance, hyperarousal, and negative alterations in cognition mood.^2^ Moreover, people with PTSD often report difficulties with concentration, attention, and memory,^2–4^ which are associated with decreased social and occupational performance and quality of life.^5,6^ Working memory (WM) is one of the cognitive processes affected in individuals with PTSD,^7^ yet the neural underpinnings of WM dysfunction are not well understood. In addition to the foundational role WM plays in daily life, WM facilitates some PTSD treatment strategies, such as cognitive behavioral therapy.^8^ A better understanding of the relationship between PTSD and WM deficiency could help inform therapeutic solutions to address the cognitive limitations of individuals with PTSD.

The brain’s response to a stressor involves reciprocal circuitry of the amygdala and prefrontal cortex (PFC).^9^ Exposure to a stressor activates the amygdala, which engages prefrontal glutamatergic circuits involved in regulating emotional and behavioral responses. In turn, glutamatergic projections from the PFC activate GABAergic interneurons within the amygdala, providing inhibitory control over amygdala activity. In PTSD, this regulatory circuit appears to be disrupted, with hyperactivity of the amygdala accompanied by hypoactivity of prefrontal regions,^10^ including the dorsolateral prefrontal cortex (DLPFC). The DLPFC plays a central role in WM,^10–12^ and functional neuroimaging studies have reported altered DLPFC activation during WM tasks in individuals with PTSD.^10,13,14^ Despite the established relationship between PTSD and WM impairment, the neurobiological mechanisms linking altered DLPFC function to cognitive dysfunction remain poorly understood.

The glutamatergic system has been implicated in the pathophysiology of PTSD.^9,15–18^ Animal models suggest that processes of learned fear rely on glutamatergic and GABAergic mechanisms in various brain regions, including the PFC.^19,20^ Further studies suggest that stress increases glutamate release in general^21–23^ and glutamate efflux rate in the PFC.^24^ In addition, glutamate levels appear to correlate with symptom severity as well as cognitive performance measures like learning and memory.^25–28^ Lastly, investigations involving the administration of drugs that enhance glutamatergic function have found improvements in PTSD symptoms.^29–35^ Altogether, these studies indicate a relationship between PTSD and glutamatergic function.

In addition to its relevance to PTSD, glutamatergic function has been associated with WM performance in the PFC and DLPFC. Neuroimaging studies have found the DLPFC to be consistently activated in WM tasks.^10^ Although studies examining DLPFC activation during WM tasks in PTSD have reported mixed findings,^10,36–38^ possibly reflecting differences in task design and study populations, these findings collectively suggest that WM-related neural processes are disrupted in PTSD, potentially due to elevated levels of glucocorticoids.^39^ Furthermore, WM appears to be particularly susceptible to the effects of glucocorticoids compared with declarative memory,^40^ suggesting the PFC is especially vulnerable to stress. Together, these findings suggest that glutamatergic function within the DLPFC may play an important role in WM dysfunction associated with PTSD.

Magnetic resonance spectroscopy (MRS) studies have provided evidence for impaired neurochemical systems in PTSD.^41^ MRS can measure neurometabolite levels *in vivo* and non-invasively. In the past, lower field strength scanners have limited the study of metabolites with significant spectral overlap, such as glutamate and glutamine. MRS at higher field strengths, such as 7T, allows the separation of these overlapping signals. Despite these developments in MRS technology, there is a lack of MRS studies of the DLPFC in individuals with PTSD.^41^ Recently, using 7T MRS, we found lower glutamate in the DLPFC of trauma-exposed individuals with and without PTSD compared to people without trauma exposure.^42^ Another recent 7T study reported a lower prefrontal GABA-to-glutamine ratio in individuals with PTSD.^43^ Together, these findings extend earlier MRS studies performed primarily at lower field strengths, which reported lower glutamate, glutamine, or Glx (glutamate + glutamine) in regions of the PFC of trauma-exposed individuals^44,45^ and those with PTSD.^46,47^

Functional MRS (fMRS) is an advanced MRS technique that measures dynamic changes in neurometabolite concentrations during functional activation. Previous fMRS studies have reported task-related glutamate changes in response to sensory stimulation as well as during emotional and cognitive paradigms, including WM paradigms.^48–50^ Although WM-related glutamate responses have been examined in healthy individuals and several clinical populations,^51–56^ to our knowledge, no studies have investigated task-related glutamate dynamics in PTSD. Characterizing glutamatergic responses during WM may provide additional insight into the relationship among DLPFC function, glutamatergic regulation, and PTSD beyond that afforded by resting-state MRS alone.

Given the evidence for PFC dysfunction and glutamatergic alterations in individuals with PTSD, we hypothesized that reduced glutamate in PTSD would be associated with impaired WM. To test this hypothesis, we used 7T fMRS to measure dynamic glutamate levels in the DLPFC during a WM task. By characterizing glutamatergic responses during WM, the findings of the present study using fMRS extend previous resting-state MRS findings and thereby improve our understanding of neurochemical alterations associated with trauma and PTSD.

## 2. METHODS

### 2.1 Participants

The Auburn University Institutional Review Board approved this study. All participants provided written informed consent. The study sample included 80 participants aged 19 to 55 as previously reported.^42^ Participants completed the Life Events Checklist for *DSM-5* extended version (LEC-5), PTSD Checklist for *DSM-5* (PCL-5), Beck Depression Inventory (BDI-II), and Beck Anxiety Inventory (BAI). Participants were assigned one of three groups as previously described^42^: PTSD (*n* = 27), trauma-exposed without PTSD (TE, *n* = 27), or no trauma exposure (NT, *n* = 26). Exclusion criteria were neurological disorders, history of concussion with loss of consciousness, alcohol use disorder, substance use disorder, psychosis and schizophrenia spectrum disorders, bipolar and related disorders, obsessive-compulsive disorder, personality disorders, and autism spectrum disorder.

### 2.2 N-Back Task

The task was a letter n-back WM task in a block design with 0-back, 1-back, and 2-back conditions.^57,58^ E-Prime (version 3.0) and a fiber optic response pad (Current Designs) were used to present the stimuli and record participant responses and reaction times. The stimuli were uppercase consonants presented for 500 ms followed by 2,000 ms of interstimulus interval (blank screen). In the 0-back condition, participants pressed a button when the letter ‘X’ appeared on the screen. In the 1-back condition, participants pressed a button when the current letter matched the preceding letter. In the 2-back condition, participants pressed a button when the current letter matched the letter presented two trials earlier (Figure 1). A 10-second instruction slide was shown at the beginning of each block. Participants completed 3 blocks each of the 0-back, 1-back, and 2-back conditions in a fixed order across all participants (Figure 1). Each block included 32 trials with 10 targets. The n-back task was followed by a fixation block consisting of a 10-second instruction slide and an 80-second fixation cross to assess recovery toward baseline glutamate levels after the task.

**Figure 1.**
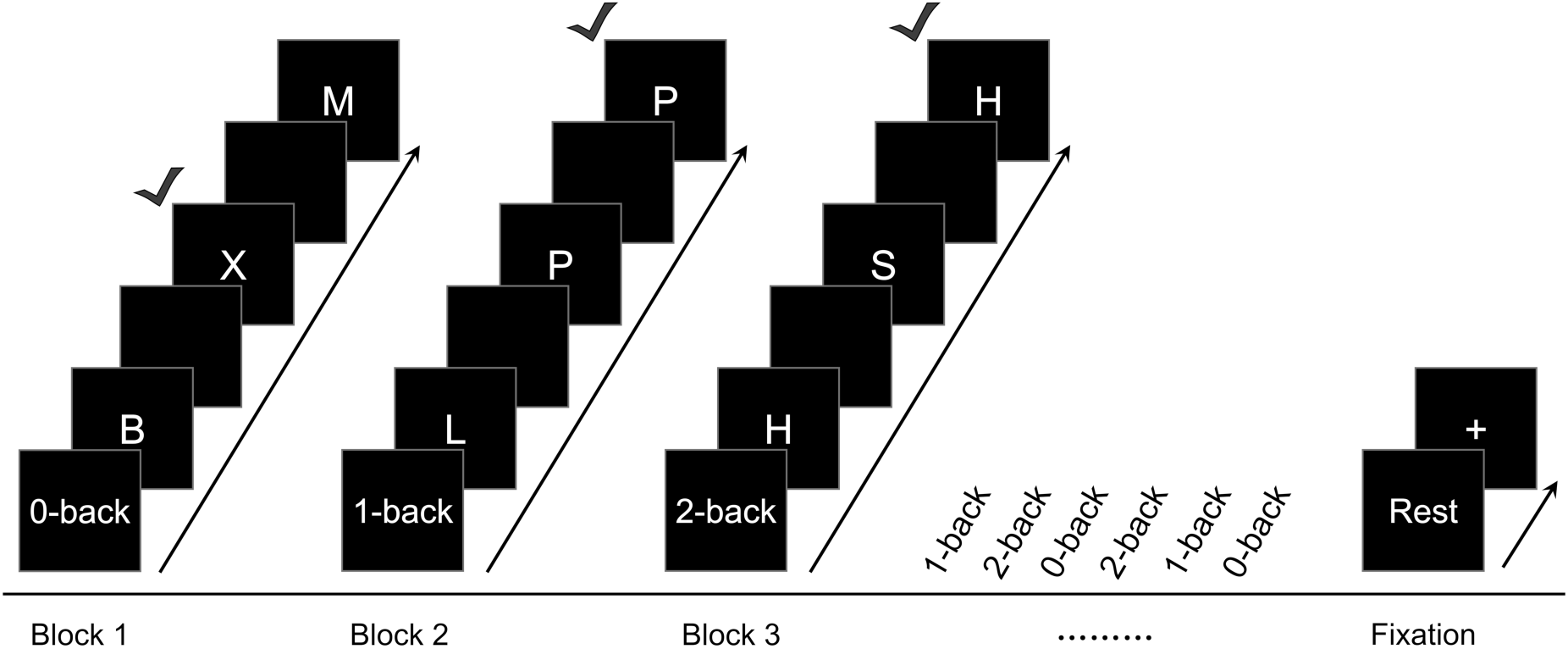
N-back task. Each block was 90 seconds long (10 seconds of instructions, 80 seconds of trials) with 32 trials and 10 targets. Each trial was 0.5 seconds of stimulus and 2 seconds of inter-stimulus interval (blank screen). The fixation block was 10 seconds of instructions and 80 seconds of crosshair.

Prior to scanning, participants completed a practice version of the n-back task outside the scanner, consisting of 1 block each of the 0-back, 1-back, and 2-back conditions with 20 trials and 5 targets per condition. Participants were required to correctly identify at least 12 of the 15 target trials (80% overall accuracy) prior to the MRI scan.

### 2.3 MRS Data Acquisition

Data were acquired on a Siemens 7T MAGNETOM scanner with a single-channel transmit and 32-channel receive head coil (Nova Medical). Structural images were acquired for anatomical reference and MRS voxel segmentation (MPRAGE, TR/TE/TI = 2200/2.82/1050 ms, flip angle = 7°, field of view = 256 × 256 mm, matrix = 256 × 256, slice thickness = 1 mm, gap = 0.5 mm, 192 slices, GRAPPA acceleration factor = 2, sagittal acquisition). The MRS voxel was placed in the left DLPFC parallel to the cortical surface (Figure 2). A 25 × 25 × 25 mm^3^ voxel was used to provide increased signal-to-noise ratio (SNR) for the functional scans. Following FASTESTMAP shimming and voxel-based flip angle calibration, spectra were acquired using an ultra-short TE STEAM sequence (TR/TE/TM = 10,000/5/45 ms, 4 kHz spectral bandwidth, 2048 points) with outer volume suppression and VAPOR water suppression^59–62^ (Table S1).

**Figure 2.**
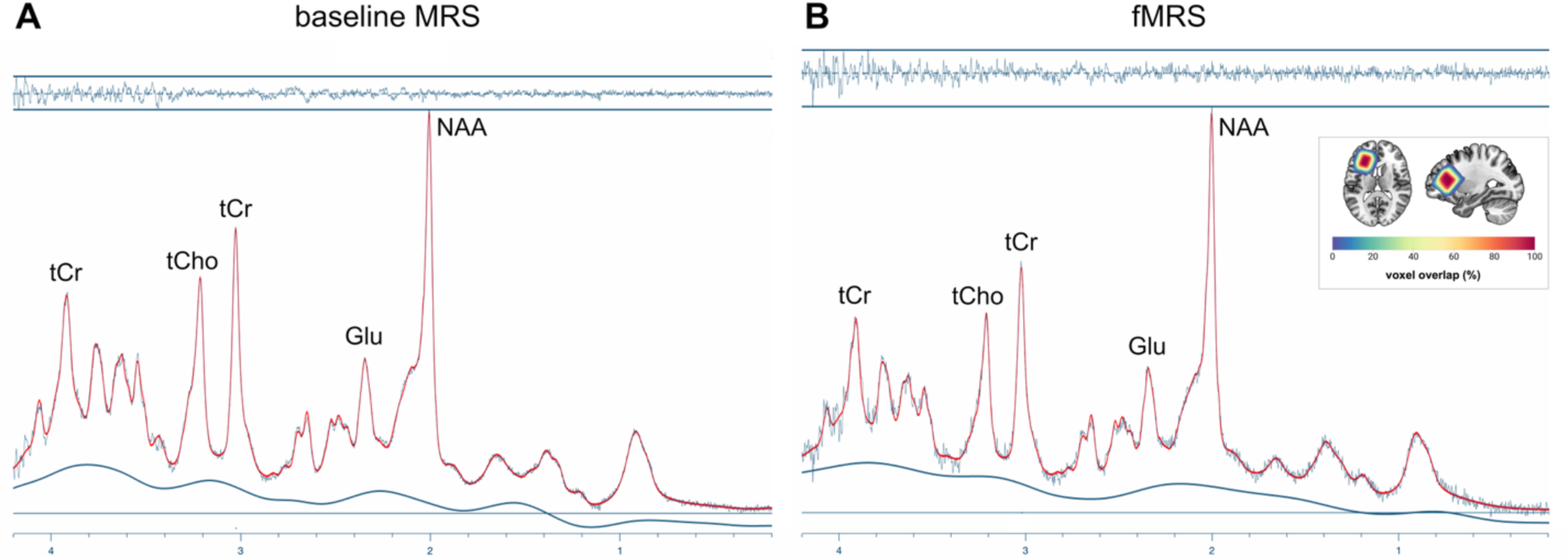
Representative spectra. (A) The baseline spectrum was acquired with 32 averages. (B) The spectrum for each fMRS block comprised 8 averages. (Inset) Spectra were acquired from a voxel in the left dorsolateral prefrontal cortex (25 ◻ 25 ◻ 25 mm2). Color scale indicates the voxel overlap across participants. Abbreviations: Glu, glutamate; NAA, N-acetylaspartate; tCho, total choline (glycerophosphocholine + phosphocholine); tCr, total creatine (creatine + phosphocreatine).

Prior to the fMRS scan, a baseline MRS scan was performed to measure resting glutamate concentrations as previously reported.^42^ Participants were instructed to focus on a crosshair during the baseline scan, and 32 water-suppressed FIDs were acquired and averaged for each participant. During each fMRS block, 9 FIDs were acquired, comprising 1 FID during the instruction slide and 8 FIDs during the task. Spectra without water suppression (4 FIDs) were acquired after the baseline MRS scan and after the fMRS scan.

### 2.4 Data Processing

#### 2.4.1 Baseline MRS

Coil-combined and averaged spectra from the baseline scans were exported as a single DICOM file for each participant and processed in Matlab (Mathworks 2024b) using the Osprey toolbox (2.9.6).^63^ Baseline spectra were preprocessed in Osprey, including removal of the residual water signal and eddy current correction, and were fit with LCModel^64^ (verion 6.3-1N implemented in Osprey) in the range of 0.2-4.2 ppm using a simulated basis set^42^ (Table S1). Tissue within the MRS voxel was semented into gray matter, white matter, and cerebrospinal fluid using SPM12. Tissue– and relaxation-corrected glutamate concentrations were quantified using Osprey. Spectra were excluded if the creatine (Cr) linewidth was greater than 30 Hz. Glutamate concentrations were excluded from statistical analysis if the Cramer-Rao lower bound (CRLB) exceeded 20%.

We previously reported baseline glutamate concentrations for this sample.^42^ For transparency, we note that these baseline spectra were reanalyzed following an update to Osprey that modified the metabolite-specific relaxation times. While this resulted in differently scaled glutamate concentrations, the underlying data distributions were unaffected (Figure S1).

#### 2.4.2 Functional MRS

The coil-combined individual FIDs (one DICOM file per FID) from the functional scans were sorted by block. The FID acquired during the instruction slide was excluded prior to preprocessing so that each functional spectrum represented the 8 FIDs acquired during task performance. The 8 task FIDs within each block were preprocessed in Osprey using probabilistic spectral alignment and were fit with LCModel as described above. Each participant had 10 functional spectra corresponding to 9 n-back blocks and 1 post-task fixation block. Tissue– and relaxation-corrected glutamate concentrations were quantified for each block. The same exclusion criteria as above were applied.

#### 2.4.3 Behavioral Measures

N-back task performance was assessed using percent correct, reaction time (RT), and the discrimination index (*d’*).^58,65,66^ Participants’ responses were classified as hits (i.e., the participant pushed the button when the stimulus was a target), false alarms (FA; i.e., the participant pushed the button when the stimulus was not a target), and misses (i.e., the participant did not push the button when the stimulus was a target). The hit rate [HR = (0.5 + hits)/(number of targets + 1.0] and false alarm rate [FAR = (0.5 + FA)/(1 + (1 – number of targets))] were then calculated and used to compute the discrimination index (*d’* = *Z*_FAR_ – *Z*_HR_).^66,67^ Higher *d’* values indicate a greater ability to discriminate between targets and non-targets. Percent correct was calculated as the percentage of correctly classified trials (targets and non-targets) across the three blocks within each WM load.

### 2.5 Statistical Analysis

Statistical analyses were conducted in R/RStudio (version 4.4.1). Statistical significance was set at two-tailed *p* < 0.05. Demographics and clinical measures were analyzed as previously reported.^42^

Behavioral metcis (percent correct, reaction time, *d’*) were analyzed using a repeated measures ANOVA with group (PTSD, TE, NT) as the between-subject factor and WM load (0-back, 1-back, 2-back) as the within-subject factor. Greenhouse-Geisser correction was applied to within-subject effects and interactions when the assumption of sphericity was violated. Significant effects and interactions were followed by Tukey-adjusted pairwise comparisons of estimated marginal means.

Participant-level glutamate values were excluded if they were more than three median absolute deviations from the median glutamate value across all groups within a given n-back condition.^68^ Glutamate concentrations were analyzed using a linear mixed-effects model with fixed effects of group (PTSD, TE, NT), condition (baseline, 0-back, 1-back, 2-back, fixation), and their interaction. Participant was included as a random intercept to account for repeated measurements. Age and sex were included as covariates. Spectral quality metrics (SNR, linewidth, CRLB) were analyzed using linear mixed-effects models with fixed effects of group and condition and a random intercept for participant. Significant main effects and interactions were followed by Tukey-adjusted pairwise comparisons of estimated marginal means.

## 3. RESULTS

### 3.1 Participants

The sample included 27 individuals with PTSD, 27 individuals who were exposed to trauma but did not have PTSD (TE), and 26 individuals with no trauma exposure (NT) as previously described.^42^ Demographic and clinical characteristics are summarized in Table 1. Age, sex, race, and ethnicity did not differ significantly among the groups. One participant in the PTSD group completed only the baseline MRS scan and was excluded from the behavioral analyses and task-related fMRS analyses.

**Table 1.** Demographics and Clinical Characteristics.

|  | PTSD | TE | NT | Statistic | *p* |
| --- | --- | --- | --- | --- | --- |
| Sample size, *n* | 27 | 27 | 26 |  |  |
| Age, mean (SD) | 27.85 (8.32) | 31.56 (9.20) | 29.08 (10.97) | $F(2,77) = 1.06$ | 0.35 |
| Sex, *n* | | | | $\chi^2(2) = 4.01$ | 0.14 |
| Female | 21 | 14 | 16 |  |  |
| Male | 6 | 13 | 10 |  |  |
| Race, *n* |  |  |  | Fisher's exact | 0.55 |
| Asian | 1 | 1 | 4 |  |  |
| Multiracial | 2 | 2 | 1 |  |  |
| White | 24 | 24 | 21 |  |  |
| Ethnicity, *n* |  |  |  | Fisher's exact | 0.87 |
| Hispanic or Latino | 2 | 3 | 1 |  |  |
| Non-Hispanic or Non-Latino | 25 | 24 | 25 |  |  |
| BDI, mean (SD) | 26.37 (10.16)<sup>a,b</sup> | 7.96 (6.14) | 4.81 (6.00) | $F(2,77) = 61.24$ | < 0.001 |
| BAI, mean (SD) | 25.41 (9.94)<sup>a,b</sup> | 8.15 (6.83)<sup>c</sup> | 2.65 (3.26) | $F(2,77) = 71.58$ | < 0.001 |
| PCL-5, mean (SD) | 50.93 (11.59) | 11.52 (9.63) | --- | $t(52) = 13.59$ | < 0.001 |

Abbreviations: BAI, Beck Anxiety Inventory; BDI, Beck Depression Inventory; NT, no trauma exposure; PCL-5, PTSD Checklist for *DSM-5*; PTSD, posttraumatic stress disorder; TE, trauma-exposed without PTSD
<sup>a</sup>Significant difference between PTSD and TE: $p_{\text{Tukey}} < 0.001$ .
<sup>b</sup>Significant difference between PTSD and NT: $p_{\text{Tukey}} < 0.001$ .
<sup>c</sup>Significant difference between TE and NT: $p_{\text{Tukey}} = 0.02$ .

### 3.2 N-back Task Performance

Results of the n-back behavioral performance are shown in Tables 2 and 3. Percent correct showed significant main effects of group and WM load. While percent correct tended to be lower in the PTSD group compared to the TE and NT groups, no significant differences were observed between groups (PTSD vs. TE: *p*_Tukey_ = 0.06; PTSD vs. NT: *p*_Tukey_ = 0.10; TE vs. NT: *p*_Tukey_ = 0.97). Percent correct decreased with increasing WM load, with significantly lower accuracy during the 2-back condition than the 0-back (*p*_Tukey_ < 0.001) and 1-back (*p*_Tukey_ < 0.001) conditions and during the 1-back condition than the 0-back condition (*p*_Tukey_ < 0.001).

**Table 2.** N-back Task Performance (mean (SD))

| Measure | Load | PTSD | TE | NT |
| --- | --- | --- | --- | --- |
| Percentage Correct | 0-back | 99.36 (1.64) | 99.61 (1.09) | 99.68 (0.77) |
|  | 1-back | 98.00 (2.65) | 98.96 (1.12) | 99.48 (0.68) |
|  | 2-back | 95.11 (4.43) | 97.30 (2.11) | 96.39 (3.80) |
| Reaction Time (ms) | 0-back | 415.67 (72.29) | 416.96 (72.41) | 403.15 (55.83) |
|  | 1-back | 455.52 (102.05) | 446.87 (110.26) | 422.83 (78.74) |
|  | 2-back | 510.50 (135.37) | 500.24 (137.13) | 476.62 (111.35) |
| d' | 0-back | 4.37 (0.41) | 4.46 (0.29) | 4.46 (0.25) |
|  | 1-back | 3.93 (0.58) | 4.16 (0.41) | 4.35 (0.29) |
|  | 2-back | 3.44 (0.64) | 3.78 (0.48) | 3.67 (0.74) |

Abbreviations: NT, no trauma exposure; PTSD, posttraumatic stress disorder; TE, trauma-exposed without PTSD

**Table 3.** Repeated-Measures ANOVA Results for N-back Task Performance.

| Measure | Effect | *F* | *p* |
| --- | --- | --- | --- |
| Percentage Correct | Group | $F(2,76) = 3.20$ | 0.046 |
| | Load | $F(1.46,110.79) = 62.99$ | < 0.001 |
| | Group × Load | $F(2.92,110.79) = 2.22$ | 0.092 |
| Reaction Time | Group | $F(2,76) = 0.60$ | 0.552 |
| | Load | $F(1.42,107.63) = 50.27$ | < 0.001 |
| | Group × Load | $F(2.83,107.63) = 0.33$ | 0.792 |
| $d'$ | Group | $F(2,76) = 3.60$ | 0.032 |
| | Load | $F(1.78,135.64) = 89.23$ | < 0.001 |
| | Group × Load | $F(3.57,135.64) = 1.94$ | 0.116 |

RT showed a significant main effect of WM load but no significant effect of group. RT increased with increasing WM load, with significantly slower responses during the 2-back condition than the 0-back (*p*_Tukey_ < 0.001) and 1-back (*p*_Tukey_ < 0.001) conditions and during the 1-back condition than the 0-back condition (*p*_Tukey_ < 0.001).

The discrimination index (*d’*) showed significant main effects of group and WM load. The PTSD group had lower *d’* values than the NT group (*p*_Tukey_ = 0.04), whereas the TE group did not differ from either the PTSD (*p*_Tukey_ = 0.08) or NT (*p*_Tukey_ = 0.95) groups. The discrimination index decreased with increasing WM load, with significantly lower *d’* values during the 2-back condition than the 0-back (*p*_Tukey_ < 0.001) and 1-back (*p*_Tukey_ < 0.001) conditions and during the 1-back condition than the 0-back condition (*p*_Tukey_ < 0.001).

### 3.3 Glutamate

A total of 870 spectra were acquired across all participants. Based on the predefined quality criteria, 42 spectra were excluded. Following quality control, 17 additional spectra were identified as glutamate outliers and excluded from subsequent analyses. Representative baseline and fMRS spectra are shown in Figure 2.

Glutamate concentrations are summarized in Table 4 and Figure 3, and linear mixed-effects model results are presented in Table 5. Significant effects of group, condition, and group × condition interaction were observed. Age and sex were also significant covariates. Glutamate was lower in the PTSD group than the NT group at baseline (*p*_Tukey_ = 0.03), 0-back (*p*_Tukey_ = 0.02), 1-back (*p*_Tukey_ < 0.001), and 2-back (*p*_Tukey_ < 0.001). The TE group showed lower glutamate than the NT group during the 1-back (*p*_Tukey_ = 0.005) and 2-back (*p*_Tukey_ = 0.02) conditions. No significant group differences were observed during the fixation condition. Within the NT group, glutamate was significantly lower at baseline than during the 0-back (*p*_Tukey_ = 0.04), 1-back (*p*_Tukey_ = 0.002), and 2-back (*p*_Tukey_ = 0.002) conditions. Glutamate during fixation was also significantly lower than during the 0-back (*p*_Tukey_ = 0.01), 1-back (*p*_Tukey_ < 0.001), and 2-back (*p*_Tukey_ < 0.001) conditions. Within the PTSD and TE groups, glutamate was significantly lower at baseline than during the 0-back condition (*p*_Tukey_ = 0.01 and 0.05, respectively).

**Figure 3.**
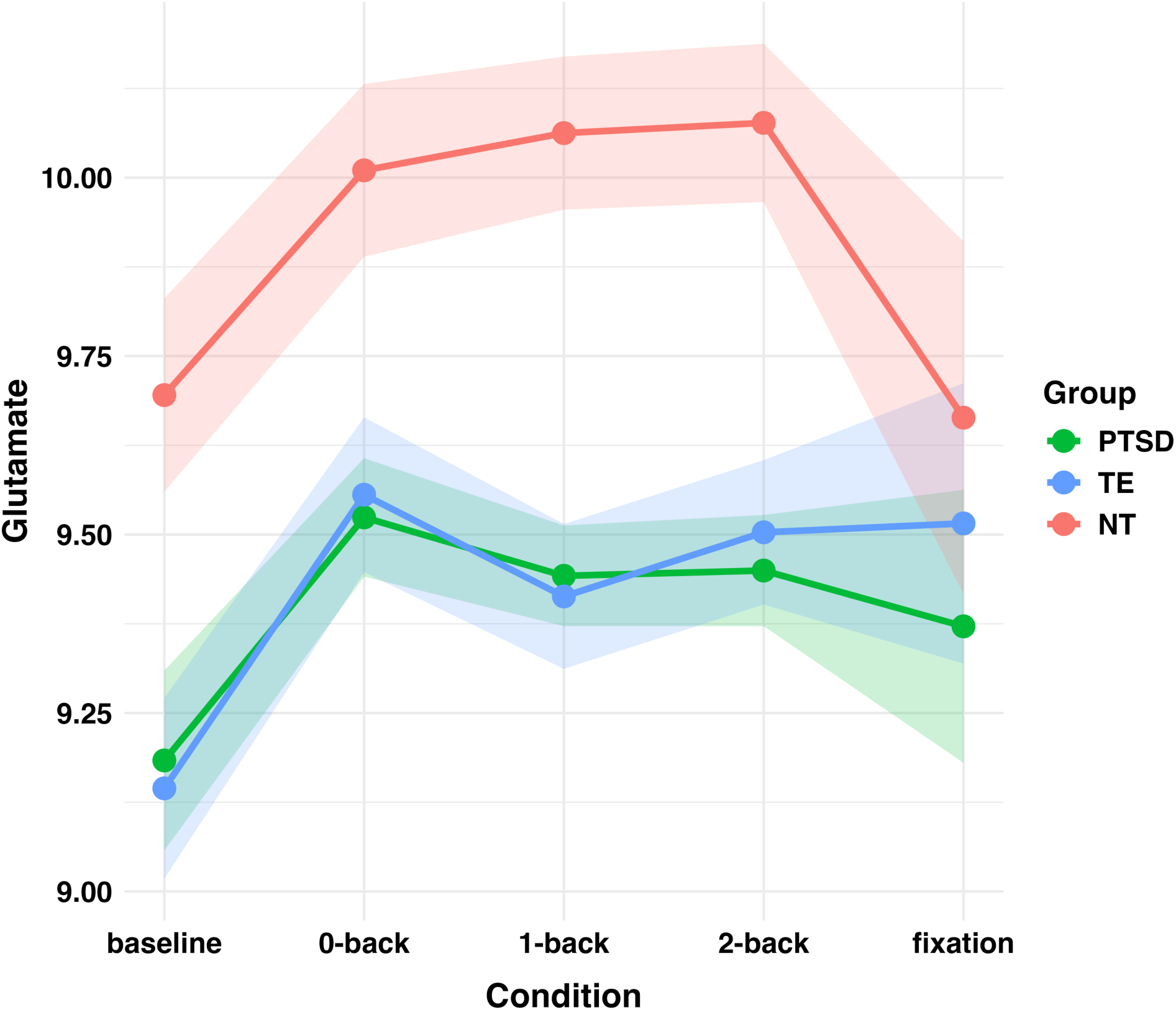
Glutamate concentrations (mean ± SEM) at baseline fixation, during the n-back task (0-back, 1-back, 2-back), and during the post-task fixation. Abbreviations: NT, no trauma exposure; PTSD, post-traumatic stress disorder; TE, trauma exposed without PTSD.

**Table 4.**
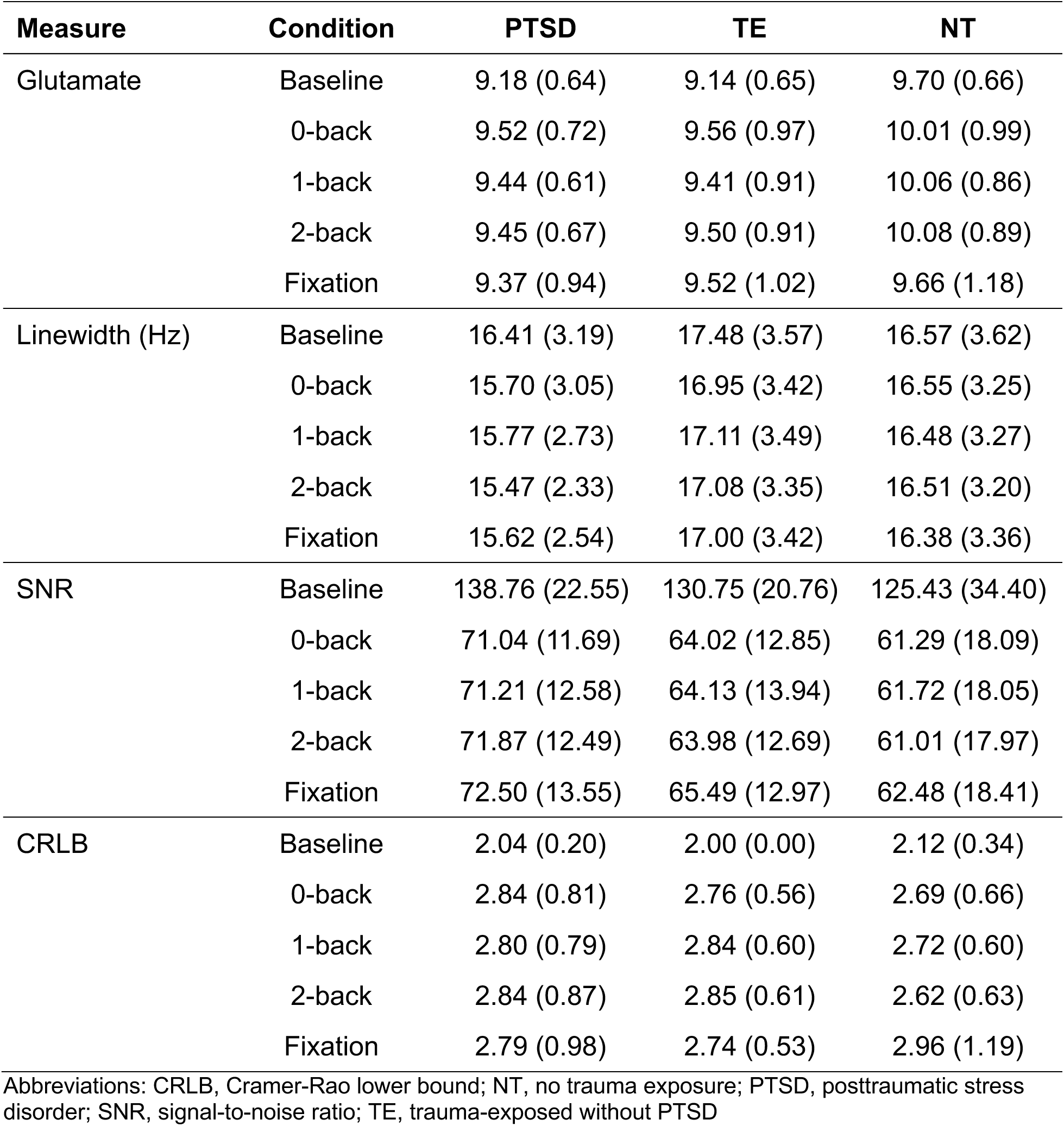
Glutamate Concentrations and Spectral Quality Metrics (mean (SD))

**Table 5.** Linear Mixed-Effects Model Results for Glutamate Concentrations and Spectral Quality Measures.

| Measure | Effect | *F* | *p* |
| --- | --- | --- | --- |
| Glutamate | Group | $F(2,73.39) = 5.33$ | 0.007 |
| | Condition | $F(4,724.23) = 8.72$ | < 0.001 |
| | Group × Condition | $F(8,724.19) = 2.16$ | 0.03 |
| | Age | $F(1,70.97) = 15.67$ | < 0.001 |
| | Sex | $F(1,70.84) = 5.02$ | 0.03 |
| Linewidth | Group | $F(2,71.56) = 1.19$ | 0.31 |
| | Condition | $F(4,728.74) = 0.77$ | 0.54 |
| SNR | Group | $F(2,72.29) = 3.05$ | 0.053 |
| | Condition | $F(4,729.38) = 1960.86$ | < 0.001 |
| CRLB | Group | $F(2,73.48) = 0.23$ | 0.79 |
| | Condition | $F(4,732.64) = 43.51$ | < 0.001 |

Abbreviations: CRLB, Cramer-Rao lower bound; SNR, signal-to-noise ratio

Spectral quality metrics are summarized in Tables 4 and 5. SNR showed a significant effect of condition but no significant effect of group. As expected, SNR was significantly higher for the baseline spectra than for the task and fixation spectra (all *p*_Tukey_ < 0.001), whereas no differences were observed among the task and fixation spectra (*p*_Tukey_ = 0.60-0.99). Linewidth did not differ significantly by group or condition. CRLB showed a significant effect of condition but no significant effect of group. CRLB was significantly lower for the baseline spectra than for the task and fixation spectra (all *p*_Tukey_ < 0.001), with no significant differences among the task and fixation spectra (*p*_Tukey_ = 0.83-0.99).

Because baseline spectra were acquired with a greater number of signal averages than the functional spectra, an additional linear mixed-effects model was performed using only the task and fixation conditions. The pattern of findings remained unchanged. Specifically, significant effects of group, condition, and group × condition interaction were again observed (Table S2). Pairwise comparisons indicated lower glutamate in the PTSD group than the NT group in the 0-back (*p*_Tukey_ = 0.01), 1-back (*p*_Tukey_ < 0.001), and 2-back (*p*_Tukey_ < 0.001) conditions. The TE group also showed lower glutamate than the NT group during the 1-back (*p*_Tukey_ = 0.005) and 2-back (*p*_Tukey_ = 0.02) conditions. No group differences were observed during the fixation condition. Within the NT group, glutamate was lower during fixation than during the 0-back (*p*_Tukey_ = 0.01), 1-back (*p*_Tukey_ < 0.001), and 2-back (*p*_Tukey_ < 0.001) conditions. No significant differences were observed among the task conditions within any group.

To determine whether the repeated presentation of the WM conditions influenced the primary findings,^48,69^ an additional linear mixed-effects model including block as a fixed effect was performed. Inclusion of block did not significantly improve model fit (likelihood ratio test, *χ*^2^(6) = 2.46, *p* = 0.87), and the pattern of findings remained unchanged (Table S3).

## 4. DISCUSSION

In this study, we used 7T fMRS to examine glutamate changes in the DLPFC during an n-back WM task in individuals with and without trauma exposure and PTSD. Glutamate concentrations differed by group and task condition, with lower glutamate in individuals with PTSD than those with no trauma exposure (NT) at baseline and during the WM task. Individuals exposed to trauma without PTSD (TE) also exhibited lower glutamate than NT during the 1-back and 2-back conditions. In the NT group, glutamate was higher during the 0-back, 1-back, and 2-back conditions than at baseline, whereas PTSD and TE exhibited more limited task-related increases in glutamate. Behaviorally, accuracy decreased and RT increased with increasing WM load, while the discrimination index (*d’*) was lower in PTSD than NT.

Our findings support the growing evidence of altered glutamatergic dysfunction in PTSD.^9,15,16,46,47,70^ Consistent with our previous report on this sample,^42^ individuals with PTSD exhibited lower glutamate in the DLPFC at baseline compared to NT. Importantly, these differences persisted during the WM task, indicating that reduced glutamate was not limited to the resting state. Previous MRS studies of PTSD have reported altered glutamate in the prefrontal cortex and other brain regions,^41,46,47,70^ although findings have varied across studies because of differences in brain region, acquisition methods, field strength, and participant characteristics. Together, these findings suggest that altered glutamatergic function is a feature of PTSD that extends beyond resting neurochemistry and remains evident during task performance.

Individuals exposed to trauma without PTSD also exhibited lower glutamate than the NT group, although we observed these differences only during the 1-back and 2-back conditions. This pattern suggests that trauma exposure alone may influence glutamatergic responses during periods of increased cognitive demand (e.g., higher WM load), even in the absence of PTSD. Although the TE group did not differ significantly from the PTSD group, the absence of significant baseline differences and the emergence of differences only during the more demanding task conditions may indicate that functional challenges are more sensitive than resting measurements for detecting subtle neurochemical alterations associated with trauma exposure. Additional studies are needed to determine whether these findings reflect persistent neurobiological effects of trauma, compensatory adaptations, or factors associated with resilience to PTSD.

The inclusion of both baseline and functional MRS measurements provides additional insight beyond conventional resting-state MRS studies. Whereas resting-state MRS characterizes static metabolite concentrations, fMRS allows glutamate to be examined during active cognitive processing.^71^ In the present study, group differences persisted during the WM task, and the TE group also demonstrated altered glutamate during the more demanding WM loads. These findings suggest that fMRS may provide complementary information regarding glutamatergic dysfunction that is not fully captured by resting-state measurements alone and highlight the potential value of examining metabolite dynamics during cognitive engagement.

Functional MRS provides a unique opportunity to examine dynamic changes in glutamate during cognitive engagement. Previous fMRS studies have reported task-related increases in glutamate during cognitive tasks, suggesting that glutamate concentrations dynamically respond to increased neuronal demand. Although the precise physiological mechanisms underlying task-related glutamate changes remain an active area of investigation, increases in glutamate measured with fMRS are generally thought to reflect increased excitatory neurotransmission and the associated metabolic processes required to support neuronal function.^48,49,71^ In addition to metabolic changes, task-related increases in the glutamate signal may also reflect activity-dependent shifts of glutamate between compartments that differ in their visibility to MRS, resulting in changes in the measured signal without necessarily altering total tissue glutamate concentrations.^48^

Consistent with these observations, the NT group exhibited a 3.2-3.9% increase in glutamate from baseline to the WM task, followed by a 3.5-4.2% decrease during the post-task fixation period. A meta-analysis of block-design fMRS studies reported an average glutamate increase of approximately 4.7%,^48^ and one fMRS study of WM reported a 2.7% increase in DLPFC glutamate during 2-back performance.^53^ The magnitude of the glutamate increase we observed in the NT group was consistent with these estimates. Similar task-related increases followed by a gradual return toward baseline have also been reported in healthy individuals performing a Stroop task.^72^ In contrast, glutamate increased by 2.8-3.7% in the PTSD group and 3.0-4.6% in the TE group during the task. Unlike the NT group, neither the PTSD nor TE groups demonstrated significant reductions in glutamate during the post-task fixation period. Taken together, these findings are consistent with previous reports describing altered task-related glutamate modulation in clinical populations.^54,56,73^

Previous fMRS studies have reported both increases and null effects in task-related glutamate responses during WM and other cognitive paradigms. In healthy individuals, increased DLPFC glutamate has been observed during WM,^52,53^ whereas other studies have reported no detectable change in Glx,^51^ suggesting that task-related glutamatergic responses may depend on factors such as acquisition methods, temporal resolution, and experimental design. Importantly, studies of schizophrenia, major depressive disorder, bipolar affective disorder, and mild cognitive impairment have consistently reported task-related glutamate increases in healthy controls that were absent or attenuated in patient groups.^54,56,73^ Similarly, an fMRS study of acute pharmacological stress likewise demonstrated glutamate increases during a 2-back task under placebo conditions but not following acute stress exposure.^55^ Together, these studies indicate that task-related glutamate responses vary across brain regions, cognitive paradigms, and clinical populations and may be influenced by methodological, clinical, or physiological factors. The present study extends this literature by demonstrating altered patterns of glutamate modulation across resting state, multiple working memory loads, and post-task fixation in trauma exposed individuals with and without PTSD.

The present findings do not identify the biological mechanisms responsible for the altered glutamate modulation observed in the PTSD and TE groups. Although fMRS measures total tissue glutamate rather than synaptic glutamate release, several processes may contribute to these differences, including altered glutamatergic neurotransmission, impaired glutamate-glutamine cycling, altered metabolic function, or diminished recruitment of prefrontal cortical networks during WM. Future studies combining fMRS with complementary neuroimaging or pharmacological approaches may help distinguish among these possibilities.

Despite the observed neurochemical differences, behavioral differences between the groups were relatively modest. As expected, increasing WM load was associated with lower accuracy, slower reaction times, and reduced discrimination performance, consistent with previous WM studies.^58,65^ The PTSD group exhibited lower *d’* values than the NT group, indicating reduced ability to distinguish targets from non-targets. However, percent correct did not differ significantly between groups after correction for multiple comparisons, and reaction time was comparable across groups.

The relationship between trauma exposure, PTSD, and WM performance remains incompletely understood. Several studies have reported impaired WM performance in trauma-exposed individuals and those with PTSD,^10,13,14^ whereas others have observed relatively preserved performance, resulting in a lack of consensus regarding the presence and magnitude of cognitive impairments in PTSD.^7^ Discrepant findings across studies have been attributed to methodological and clinical factors, including comorbid psychiatric disorders and traumatic brain injuries,^74,75^ differences in the type and duration of trauma exposure,^76,77^ and pre-trauma factors that may influence cognitive function.^78^

The relatively modest behavioral differences observed in the present study, despite significant glutamatergic abnormalities, may suggest that neurochemical alterations are detectable even when behavioral performance is only subtly affected. One possibility is that individuals with PTSD engage compensatory neural processes to maintain performance during WM tasks, resulting in relatively preserved behavioral performance despite altered glutamatergic function. Alternatively, glutamatergic abnormalities may precede measurable behavioral impairments and therefore represent a more sensitive marker of altered prefrontal function than behavioral measures alone.

Finally, the choice of task may also have contributed to the relatively small behavioral differences observed. We used a letter n-back task to isolate WM processing without introducing trauma-related or emotionally salient stimuli. Tasks incorporating emotional or trauma-related content may more effectively reveal cognitive differences associated with PTSD and should be explored in future studies. Taken together, these findings suggest that fMRS may detect alterations in glutamatergic function that are not readily apparent from behavioral performance alone.

Our findings should be interpreted in the context of several limitations. First, glutamate was measured in a single voxel in the left DLPFC. Although this region plays a central role in WM and has been implicated in PTSD, the findings may not generalize to other brain regions involved in cognitive or affective processing. Second, we did not functionally define our voxel location for each participant.^79^ We manually prescribed our acquisition voxel for each participant to cover the DLPFC, but the voxel may not have been placed to optimally capture the regions of the DLPC most strongly engaged during the WM task. Third, baseline spectra were acquired with a greater number of signal averages than the functional spectra, resulting in higher SNR. However, repeating the primary analyses using only the task and post-task fixation spectra yielded similar results, supporting the robustness of the findings. Fourth, task-related BOLD effects can alter spectral linewidth and potentially influence metabolite quantification.^80,81^ Although some fMRS studies apply linewidth correction to minimize this effect, we did not perform such a correction. However, spectral linewidths did not differ significantly across task conditions, suggesting that task-related linewidth changes were unlikely to account for the observed glutamate findings. Fifth, the temporal resolution of the fMRS acquisition limited our ability to characterize the precise timing of glutamate responses during WM. Each functional spectrum represented the average of 8 acquisitions obtained over 80 seconds using a TR of 10,000 ms. Although previous studies suggest that task-related glutamate changes occur rapidly and can persist for approximately one minute,^72^ the present acquisition was designed to characterize sustained glutamate responses over each task block rather than rapid transient changes. Future studies using shorter TRs may provide additional insight into the temporal dynamics of glutamate during WM. Finally, although age and sex were included as covariates, other sources of clinical heterogeneity, including medication use and trauma characteristics, may have contributed to variability in glutamate concentrations and should be considered in future investigations.

In this study, we examined WM performance and task-related glutamate dynamics in individuals with PTSD, trauma exposure without PTSD, and no trauma exposure using 7T fMRS in the DLPFC. To our knowledge, this is the first fMRS study of PTSD. Consistent with previous MRS studies, individuals with PTSD exhibited lower glutamate than those without trauma exposure at baseline and throughout the WM task, whereas trauma-exposed individuals without PTSD also showed lower glutamate during the more demanding WM conditions. Within-group analyses further demonstrated dynamic glutamate modulation during working memory in the NT group, whereas these task-related changes were more limited in the trauma-exposed groups. Together, these findings provide further evidence of altered glutamatergic function associated with trauma exposure and demonstrate the potential of fMRS to characterize dynamic glutamatergic responses during cognitive processing.

## ACKNOWLEDGMENTS

We thank the participants who volunteered their time for this research. The research reported in this publication was supported by the National Institute of Mental Health of the National Institutes of Health under award number K01MH115272 (to M.A.R.).

## Abbreviations

BAI: Beck Anxiety Inventory
BDI-II: Beck Depression Inventory
CRLB: Cramer-Rao lower bound
d’: discrimination index
DLPFC: dorsolateral prefrontal cortex
FA: false alarm
FAR: false alarm rate
GABA: gamma-aminobutyric acid
HR: hit rate
LEC-5: Life Events Checklist for *DSM-5*
PCL-5: PTSD Checklist for *DSM-5*
PFC: prefrontal cortex
PTSD: post-traumatic stress disorder
RT: reaction time
WM: working memory

