## Supplemental Material for "Altered Dorsolateral Prefrontal Glutamate Dynamics During Working Memory in Trauma-Exposed Individuals With and Without PTSD: A 7T Functional Magnetic Resonance Spectroscopy Study"

for

**Table S1.** Minimum reporting standards for *in vivo* magnetic resonance spectroscopy (MRSinMRS)

| **MRSinMRS** | |
| --- | --- |
| **Site:** Auburn University MRI Research Center | |
| 1. **Hardware** | |
| a. Field strength | 7T |
| b. Manufacturer | Siemens |
| c. Model | MAGNETOM (software version VB17) |
| d. RF coil | ^1^H, single-channel transmit and 32-channel receive head coil (Nova Medical) |
| e. Additional hardware | E-Prime 3.0 running on Windows PC and a 4-button fiber optic response pad (Current Designs) |
| 2. **Baseline/Resting MRS Acquisition** | |
| a. Pulse sequence | STEAM (Siemens WIP 643) |
| b. Volume of interest (VOI) | dorsolateral prefrontal cortex |
| c. Nominal VOI size | 25 × 25 × 25 mm^3^ |
| d. Repetition time (TR), echo time (TE) | TR: 10,000 ms, TE: 5 ms |
| e. Total number of excitations or acquisitions per spectrum | 32 |
| f. Additional sequence parameters | TM: 45 ms, spectral bandwidth: 4000 Hz, number of points: 2048 |
| g. Water suppression method | VAPOR |
| h. Shimming method | FASTESTMAP followed by manual shimming of water |
| i. Triggering or motion correction method | N/A |
| 3. **fMRS Acquisition** | |
| a. Pulse sequence | STEAM (Siemens WIP 643) |
| b. Volume of interest (VOI) | dorsolateral prefrontal cortex |
| c. Nominal VOI size | 25 × 25 × 25 mm^3^ |
| d. Repetition time (TR), echo time (TE) | TR: 10,000 ms, TE: 5 ms |
| e. Total number of excitations or acquisitions per spectrum | 90 total FIDs across 10 blocks. Each block consisted of 1 FID during the instructions slide and 8 FIDs during the task. The 8 task FIDs were averaged within each block, resulting in 10 total spectra (1 spectrum per block) for each participant. |
| f. Additional sequence parameters | TM: 45 ms, spectral bandwidth: 4000 Hz, number of points: 2048 |
| g. Water suppression method | VAPOR |
| h. Shimming method | FASTESTMAP followed by manual shimming of water |
| i. Triggering or motion correction method | N/A |
| 4. **Data analysis methods and outputs** | |
| a. Analysis software | Osprey (version 2.9.6), LCModel (version 6.3-1N implemented in Osprey), SPM12 |
| b. Processing steps deviating from quoted reference or product | Baseline MRS: Spectra were coil combined and averaged by the scanner software before being exported as a single DICOM file for each participant.  fMRS: Individual FIDs were exported as DICOM files. The FID acquired during the instruction slide was excluded prior to processing so that each functional spectrum represented the 8 FIDs acquired during task performance. |
| c. Output measure | absolute concentration (approximate molal concentrations: mmol/kg of tissue water; TissCorrWaterScaled as reported by Osprey) |
| d. Quantification references and assumptions, fitting model assumptions | Fit range: 0.2-4.2 ppm  Osprey bLineKnotSpace (LCModel DKNTMN): 0.15  Simulated basis set: alanine, ascorbate, aspartate, creatine (Cr), gamma-aminobutyric acid (GABA), glucose, glutamine (Gln), glutamate (Glu), glycine, glycerophosphocholine (GPC), glutathione (GSH), lactate (Lac), myo-inositol (mI), *N*-acetylaspartate (NAA), *N*-acetylaspartylglutamate (NAAG), phosphocholine (PCh), phosphocreatine (PCr), phosphoethanolamine, scyllo-inositol, serine, taurine, and LCModel’s default macromolecules |
| 5. **Data quality** | |
| a. Reported variables | creatine SNR and creatine linewidth as reported by Osprey (Table 4) |
| b. Data exclusion criteria | creatine linewidth > 30 Hz and glutamate CRLB > 20% (Table 4) |
| c. Quality measures of postprocessing model fitting | CRLB (Table 4) |
| d. Sample spectrum | Figure 2 |

**Table S2.** Linear Mixed-Effects Model Results for Glutamate Concentrations Excluding the Baseline Condition

| **Measure** | **Effect** | ***F*** | ***p*** |
| --- | --- | --- | --- |
| Glutamate | Group | *F*(2,71.48) = 5.67 | 0.005 |
|  | Condition | *F*(3,650.83) = 3.06 | 0.03 |
|  | Group × Condition | *F*(6,650.83) = 2.77 | 0.01 |
|  | Age | *F*(1,69.63) = 13.85 | < 0.001 |
|  | Sex | *F*(1,69.55) = 5.45 | 0.02 |

**Table S3.** Linear Mixed-Effects Model Results for Glutamate Concentrations Including Block as a Fixed Effect

| **Measure** | **Effect** | ***F*** | ***p*** |
| --- | --- | --- | --- |
| Glutamate | Group | *F*(2,73.40) = 5.34 | 0.007 |
|  | Condition | *F*(4,717.89) = 5.47 | < 0.001 |
|  | Group × Condition | *F*(8,718.18) = 2.15 | 0.03 |
|  | Block | *F*(6,715.98) = 0.40 | 0.88 |
|  | Age | *F*(1,70.97) = 15.68 | < 0.001 |
|  | Sex | *F*(1,70.84) = 5.03 | 0.03 |

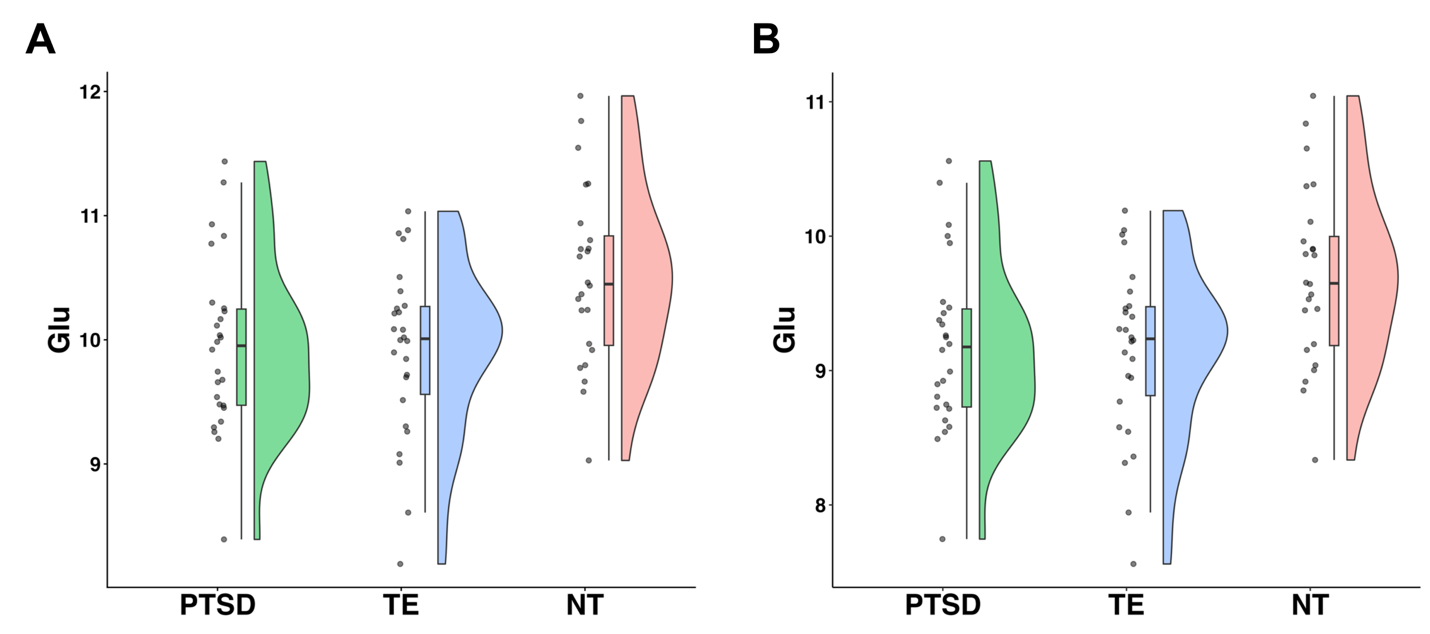

**Figure S1.** (A) Baseline resting-state glutamate concentrations for this sample were previously reported (Reid MA, Whiteman SE, Camden AA, Jeffirs SM, Weathers FW. Prefrontal metabolite alterations in individuals with posttraumatic stress disorder: a 7T magnetic resonance spectroscopy study. *Chronic Stress*. 2024;8:24705470241277451. doi:10.1177/24705470241277451). (B) The baseline spectra were reanalyzed following an update to Osprey that modified the metabolite-specific relaxation times. While this resulted in differently scaled glutamate concentrations, the underlying data distributions were unaffected.
